# Unraveling microbial interactions in the gut microbiome

**DOI:** 10.1101/2021.05.17.444446

**Authors:** Aarthi Ravikrishnan, Karthik Raman

**Affiliations:** Department of Biotechnology, Bhupat and Jyoti Mehta School of Biosciences, Indian Institute of Technology (IIT) Madras; Initiative for Biological Systems Engineering, IIT Madras; Robert Bosch Centre for Data Science and Artificial Intelligence (RBC-DSAI), IIT Madras, Chennai — 600 036, INDIA

## Abstract

The human gut comprises trillions of microorganisms that take part in several critical functions of the body. Within this structured habitat, the microbes synergise with one another mainly mediated through metabolic exchanges. In this study, we used 52 most commonly occurring microbial species in the gut and identified the interactions between them. Using our previously developed graph-based algorithm, MetQuest, we enumerate several biosynthetic pathways, mainly involved in amino acid biosynthesis and spanning across a pair of organisms. We compute the Metabolic Support Index (MSI), which captures the extent of support/interaction between a pair of microbes, based on the incremental change in metabolic capabilities achieved in a community vis-à-vis the individual organisms. Our results from these analyses are four-fold. Firstly, we show that dependencies between the gut microorganisms largely vary with respect to the environmental conditions. We observe that few phyla such as Firmicutes, Fusobacteria and Proteobacter showed significantly higher metabolic support on DMEM conditions compared to HF medium. Secondly, we infer a microbial association network based on the MSI values and found that the gut organisms are arranged in trophic-levels, with *B. bifidum* and *R. torques* acting as central nodes with the highest betweenness centrality. Thirdly, we find that species belonging to *Lactobacillus* genus, especially *L. mucosae* and *L. reuteri* show an enriched amino acid synthesising ability in the presence of many other gut organisms on a minimal glucose medium. Finally, through pathway analyses, we observe that metabolic exchanges are medium-dependent, and many of the metabolites are involved in energy metabolism, nucleotide, and vitamin biosynthesis. Overall, this study sheds light on the astonishing variety of underlying interactions between microorganisms in the gut.

**Author Summary:** Microorganisms are ubiquitous and exist in evolutionarily and metabolically diverse communities around us. In practically every ecosystem, microbes form complex dynamic assemblages—such as in the human gut, where they outnumber human cells. Community structure in these microbiomes is dictated by a complex web of interactions, where metabolic interactions are known to predominate. In this study, we employ computational tools to interrogate community metabolic networks and unravel the dependencies between microbes in the gut. We used 52 important and commonly occurring microbial species in the gut, as identified in previous studies. Through our analyses, we show that the dependencies between gut organisms vary in different environmental conditions. Further, we show that these organisms are arranged in multiple trophic levels, where a select few of them acting as central nodes. We also identify the metabolic interactions and demonstrate how synergistic interactions can enhance the amino acid biosynthesis in gut bacteria. These results, taken together, help us better understand the mechanism behind these interactions, which would pave the way for the rational design of pre- and pro-biotic formulations.

## 1 Introduction

The human gut is inhabited by trillions of micro-organisms, which carry out complex functions such as the degradation of oligosaccharides, synthesis of essential vitamins such as vitamin K and vitamin D, regulation of the immune system, and protection against pathogens [1]. Different types of micro-organisms amongst this diverse pool are responsible for performing a specific set of functions, such as the synthesis of short-chain fatty acids (SCFAs; [2]). Composition of the human gut is known to vary at different stages of life, depending on several factors such as the type of diet, host susceptibility, environmental exposure and antibiotic usage [3–7].

Within the structured habitat in the human gut, micro-organisms display complex interactions, which are often mediated through metabolic exchanges [8–10]. Microbe–microbe interactions help in maintaining diversity in the gut ecosystem. In addition, the gut micro-organisms also communicate with their human hosts through the export of metabolic by-products released from the digestion of complex polysaccharides [11]. Such interactions with the human host play a critical role in regulating several key processes, including homeostasis and inflammatory mechanisms. For instance, it was shown that SCFAs such as butyrate produced by gut bacteria serve as an energy source for colonocytes [12], help in sleep modulation [13] and induce the differentiation of immune cells [14].

Investigating the metabolism of gut microbes and how they communicate with each other and the host is thus of paramount importance. A systems-level understanding of the interactions between these gut organisms could help in gaining mechanistic insights and understanding the roles of micro-organisms. Besides, the knowledge of metabolic exchanges can also shed light on community assembly [15]. To this end, genome-scale metabolic models (GSMMs) of gut micro-organisms have emerged to be a viable tool for gaining insights into the metabolism of microbial communities. Recently, GSMMs for over 800 gut microorganisms have been reconstructed [16,17]. Several methods have been developed to analyse GSMMs [18]; these broadly fall into two major categories: constraint-based modelling [19–21] & network-based modelling [22–25]. These methods have been applied on GSMMs in a variety of studies to understand the gut metabolism under different conditions [6,17,26–29].

In this study, we seek to understand the interactions between micro-organisms in gut, by identifying several important metabolic exchanges that play critical roles in enhancing their metabolism. We used 52 important and commonly occurring microbial species in the gut, as identified in previous studies [6,26,30]. We applied our previously developed graph-based computational framework, MetQuest [25], to identify the metabolic exchanges and the associated biosynthetic pathways between 1326 pairs of organisms. We also show how amino acid biosynthesis, which has been shown to promote co-operative interactions [31], is enhanced through synergistic interactions between the gut bacteria. Further, by computing the Metabolic Support Index [32], we show that the gut organisms are arranged in multiple trophic levels, where a select few of them act as “hubs” of interaction. We also identified several metabolic exchanges between the gut organisms. These results, taken together, help us better understand the mechanism behind these interactions, which would pave the way for rational design of pre- and pro-biotic formulations.

## 2 Methods

In this section, we describe the methods we developed to carry out the analyses of the gut microbiome. Broadly, we use MetQuest [25], a method we developed previously to enumerate metabolic pathways in large metabolic networks to identify various pathways, and the metabolites exchanged therein. We also identify the metabolic dependencies between the microbes by computing Metabolic Support Index (MSI), which we developed previously [32]. We chose 52 most commonly occurring microbial species based on multiple previous studies [6,26,30] **(Supplementary Table 1)**. These organisms belong to the phyla Actinobacteria (4 species), Bacteroidetes (14 species), Firmicutes (22 species), Fusobacteria (1 species), Proteobacteria (10 species), Verrucomicrobia (1 species). We downloaded the respective genome-scale metabolic models (GSMMs) from the AGORA database (version 1.03) [16,17]. All the *in silico* analyses were carried out on three different environment conditions, i.e., minimal glucose, Dulbecco’s Modified Eagle’s medium (DMEM) and High Fibre (HF) diet conditions **(Supplementary Table 2)**, whose constituents comprised the *seed metabolites*. We added different types of mucin to the HF diet to mimic the gut conditions. All the in-house scripts used in our analyses are available on GitHub (http://github.com/aarthi31/gutmicrobiome).

### Community metabolic graph construction

We first constructed 52 individual and 1326 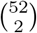 joint directed bipartite graphs using the construct_graph module in the metquest Python package (https://pypi.org/project/metquest/). Briefly, these directed bipartite graphs *G*(*M, R, E*), are constructed from the GSMMs, where metabolites (*m_i_*, ∈ *M*) and reactions (*r_j_* ∈ *R*) are the two types of nodes that are linked to each other through a directed set of edges *E*, as described in [25]. For the community metabolic network, we connect the two microbes based on their exchange reactions and assume that the two microbes exchange metabolites via a common environment [33].

### Studying microbial interactions

We studied microbial interactions by quantifying: (a) enrichment in metabolic capabilities and (b) Metabolic Support Index (MSI), using the bipartite graphs constructed. To compute these, we use outputs from the first phase of MetQuest, which include (a) scope, which captures all the metabolites that can be produced from seed set *S*, in the given metabolic network (b) reactions that are ‘stuck’, i.e., reactions whose precursor metabolites cannot be synthesised by the metabolic network using the given input conditions.

#### Enrichment in metabolic capabilities

Using the single and community metabolic graph *G* and different medium components (seed metabolites) as starting conditions, we determined the *scope* of metabolites in both these cases considered. *Scope*, as defined previously [25,34] is the set of metabolites that can be produced from the seed set *S*, in the given metabolic network. We obtained the scope and computed the increase in metabolic capabilities by comparing the scope of single and community bipartite graphs. The improvement in amino acid biosynthesis capabilities was similarly computed.

#### Metabolic Support Index

To quantify the benefits derived by the organisms in a community, we employ a metric termed Metabolic support index (MSI), previously developed by us [32]. To compute MSI, we first determined the number of reactions that are stuck, owing to the absence of precursor metabolites. MSI is defined as the fraction of blocked/stuck reactions relieved in organism *A* by the presence of another organism *B*, i.e.,

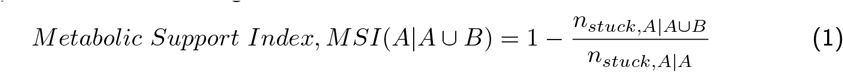

where A ∪ B denotes the community metabolic network comprising both organisms A and B, A denotes the bipartite metabolic network of organism A, *n_stuck,A|A_* denotes the number of stuck reactions in A, *n*_*stuck,A*|*A*∪*B*_ denotes the number of stuck reactions of A in A ∪ B. Further, *MSI*(*A*|*A* ∪ *B*) = 1 indicates that organism A fully benefits from the interaction with organism B, *MSI*(*A*|*A* ∪ *B*) = 0 indicates that organism A does not derive any benefit from the interaction with organism B. It is important to note that MSI values are *asymmetric*, since the two organisms (likely) have varied levels of dependencies on one another. For better clarity, we multiply the value of MSI by 100 and express it as a percentage.

Using customised Python scripts building on the guided_bfs module of MetQuest, we computed the MSI by identifying the stuck reactions in single and joint metabolic networks on two different environments/diet conditions: (i) DMEM, and (ii) HF. The MSI results were visualised using ggplot2 package [35] and the statistical analyses were carried out using inbuilt functions on R v3.6.1. Since the MSI values are asymmetric, for every pair of interactions (A→B, B→A), only the edges with highest MSI value were retained for differential network analyses, which were carried out using scripts developed in-house. Association networks were then constructed using the common set of edges that occurred in both environmental conditions. The network parameters were computed and visualised on Cytoscape v3.8.0 [36] using a hierarchical layout. Organism pairs were marked as deriving (a) equal benefits, if the absolute difference between their MSI values were less than 5%, or (b) no benefits, if the MSI values are 0.

### Computing phylogenetic and metabolic distances

The phylogenetic tree of the 52 organisms was constructed using the NCBI Taxonomy database. Briefly, the taxonomic IDs of the 52 organisms were loaded on NCBI Taxonomy Browser (http://www.ncbi.nlm.nih.gov/Taxonomy/CommonTree/wwwcmt.cgi)and the phylogenetic tree was exported as ‘PHYLIP TREE’. The phylip tree was loaded and visualised using iTOL v5 web application [37]. The annotations were suitably formatted and loaded as a separate file to be visualised on the tree. Further, to obtain the phylogenetic difference, the rooted tree (exported in ‘newick’ format from iTOL) was used to calculate the cophenetic distance using ‘ape’ package in R.

The metabolic distances between the organisms were computed using Jaccard distance, as

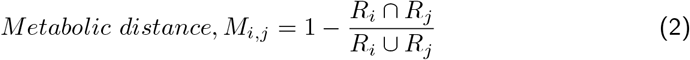

where, *R_i_* and *R_j_* are the reaction sets from the metabolic networks of organism *i* and *j* respectively *M_i,j_* value of 1 indicates that the two organisms are identical, whereas a value of 0 means that the two organisms are different. All the models were read using COBRAPy [38] and the metabolic distances were computed using in-house Python scripts.Spearman correlations between phylogenetic and metabolic distances were computed using home-grown R scripts.

### Identifying metabolic exchanges

To identify the metabolic exchanges, we first enumerated all the pathways till a pathway length cut-off of 30 using the find_pathways function in the metquest Python package on the joint bipartite graphs generated as described in the previous section. We carried out the simulations on DMEM and HF conditions. Using home-grown Python scripts, we enumerated all the pathways that spanned both the organisms, and examined each of these pathways for the presence of exchange metabolites. We then computed the unique set of metabolites by obtaining the difference between these exchange metabolite sets. We plotted these results using ‘seaborn’ library in Python.

## 3 Results

### 3.1 Metabolic dependencies vary across environment conditions

Environmental conditions often influence the interactions between different species and form the basis of their aggregations [39–41]. In order to test the interactions between the gut micro-organisms, we computed the MSI for the 1326 pairwise combinations on two different medium conditions based on their increasing richness - DMEM and HF. We observed that the MSI values on DMEM are significantly higher than that on HF conditions (one-sided Wilcoxon *p* < 0.001). This is because the number of metabolic reactions activated by the other organisms on DMEM conditions is much higher than that observed in HF conditions. Thus, the higher MSI values on DMEM conditions are in line with the expected observations.

Further, we performed a phyla-based analysis of 1326 organism combinations to understand how the metabolic interactions vary with the Phyla to which the organisms belonged. To this end, we only listed the phyla of the organisms (from the combinations) having a higher MSI value and determined how the metabolic support differed in the two conditions considered. Based on these results **(Figure 1A)**, we observed that micro-organisms belonging to the phyla Firmicutes (*p* <0.001), Fusobacter (*p* <0.001) and Proteobacter (*p* <0.01) showed significantly higher MSI values on DMEM medium compared to the HF medium (FDR corrected Wilcoxon rank-sum one-sided alternative test).

**Fig 1.**
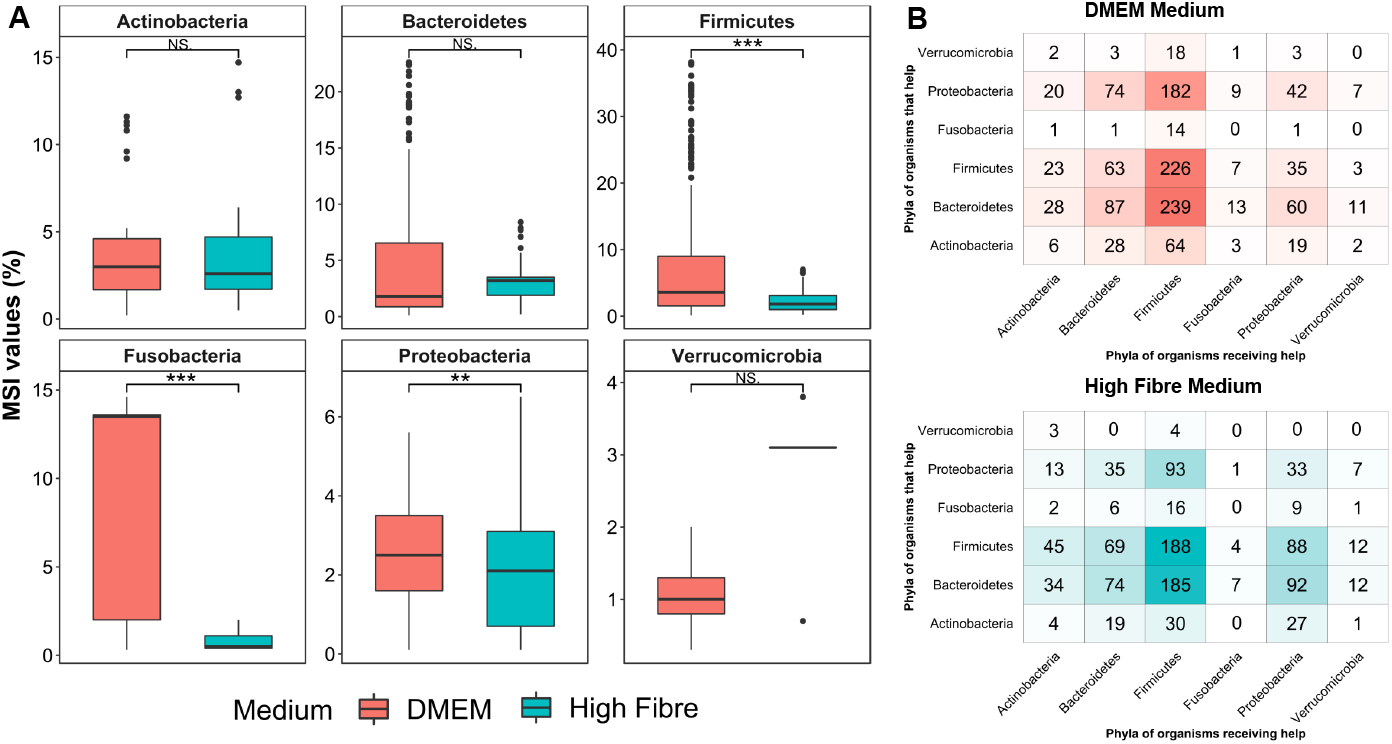
Metabolic support index (MSI) and the benefits of interactions in different environments. **(A)** MSI values grouped based on the phyla of the organism that derives the maximum benefit from the other. The values are based on two different medium conditions - (i) DMEM medium (orange colour), (ii) HF medium (blue colour). FDR-adjusted Wilcoxon one-sided p-value, ** and *** indicate *p* < 0.01 and *p* < 0.001 respectively. The number of combinations (*n*) in each category can be found in **Supplementary Table 4**. **(B)** Heatmap showing the intra- and inter-phyla interactions. The number in each cell indicates the number of combinations where a micro-organism belonging to the phyla on Y-axis helps another organism belonging to the phyla on X-axis. The top panel and the bottom panel are based on the results on DMEM and HF medium conditions respectively.

In order to check how the microbial interactions vary in different environments, we classified them into three types (see **Methods**): (i) one organism deriving higher benefit compared to the other, (ii) both organisms deriving ‘equal’ benefits, and (iii) both organisms not benefiting from one another. In total, we find that the number of microbial pairs, with one organism deriving higher benefit from the other, drops from 1295 in DMEM to 1114 in HF conditions.

From **Figure 1B**, based on the top three values from every column, we found that microorganisms belonging to Proteobacteria, Firmicutes and Bacteroidetes *help* several organisms from other phyla, irrespective of the medium conditions. It was quite interesting to note that, while the number of organisms helped by these phyla reduced in HF conditions, a few of them increased. These included the interactions between Firmicutes and Verrucomicrobia, Firmicutes and Bacteroidetes, and Firmicutes and Actinobacteria. We also observed that in the HF conditions, 203 pairs of organisms became independent, compared to only 15 in the DMEM condition. Further, the number of organism combinations that derive equal benefit from one another reduced from 16 in DMEM to 9 in HF conditions **(Supplementary Table 3)**.

### 3.2 Phylogenetic distance affects the metabolic support on different environments

Several studies have highlighted the role that phylogeny plays in deciding how the organisms aggregate together in a community [24,42,43]. Further, many studies have also pointed out that phylogeny influences species interactions in communities [44,45]. To assess the influence of phylogeny on metabolic interactions between the gut organisms, we compared the MSI values and the phylogenetic distance on DMEM and HF conditions. We observed that on DMEM, a minimal nutrient condition, the phylogenetic distance had a weak positive correlation with the MSI (Spearman rank; *ρ* = 0.22, *p* = 1.4 × 10^−15^) **(Supplementary Figure S1)**. However, no correlation was observed on HF (nutrient-rich) conditions **(Supplementary Figure S2)**. In addition, we computed the correlations between metabolic distance and phylogenetic distance to determine how it influences the MSI. As expected and shown previously [16], we also observed that the phylogenetic and the metabolic distances are positively correlated (Spearman rank; *ρ* = 0.19, *p* = 7.6 ×10^−11^) **(Supplementary Figure S3)**. However, the metabolic distance did not show any significant correlations with MSI values on both the environmental conditions. These observations imply that the microbial interactions do not depend on metabolic distance.

In addition, to determine which organism provides the most support and understand the role of phylogeny, we plotted the phylogenetic tree of the 52 organisms that we considered **(Figure 2)**. We also annotated the tree with the number of organisms each one supports, on two different medium conditions. We noted that *Bacteroides thetaiotamicron, B. uniformis, B. caccae, B. eggerthii, D. piger, Bifidobacterium longum* help many other organisms, irrespective of the medium condition considered. As described in the results §3.1 and previous studies [46], we observe that the phylum Bacteroidetes tend to *support* other organisms in the gut **(Supplementary Table 4)**. Surprisingly, a few organisms such as *Alistipes putredinis, Eubacterium rectale* and *Roseburia intestinalis* showed better support in HF conditions compared to DMEM. This points towards an enrichment in the respective metabolisms on a nutrient-rich medium, which in turn leads to better support offered by these organisms.

**Fig 2.**
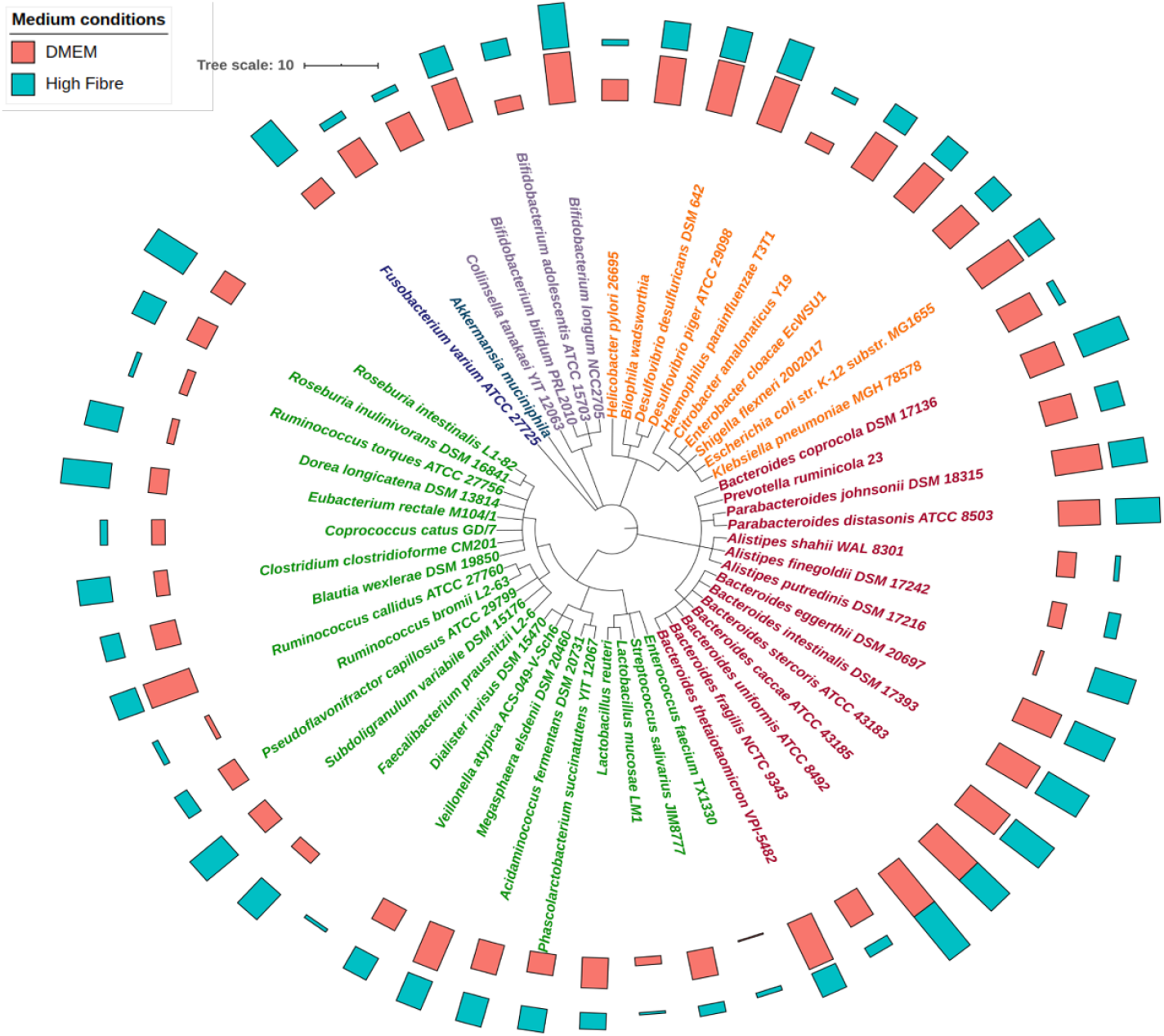
Phylogenetic tree showing the support offered by every organism. Phylogenetic tree drawn using iTOL v5 showing the relationship between 52 organisms annotated with the number of organisms supported by each of them. The two colours represent DMEM and HF medium conditions. The organisms in the tree are also coloured based on their phylum. Tree is scaled to 10 units for visulisation purposes. The outer bars (blue colour) and the inner bars (orange colour) show the number of organisms supported by each of them on HF and DMEM conditions respectively.

### 3.3 Association network suggests a trophic-level assembly of gut microbes

To further understand how the interactions between the organisms varied on different environments, we constructed an “association network” with directed edges – each edge pointing to the organism that derives maximum benefit from the other. Out of the 1295 and 1114 associations on DMEM and HF conditions respectively, we observed that 793 pairs of associations between the organisms are retained, irrespective of the environment conditions (Figure 3). From the network, we observed that, irrespective of the medium conditions, *B. uniformis* and few organisms “help” several others in the gut, pointing towards the commensalism, as shown in the previous studies [47]. In addition, we also identified a few pairs of organisms exhibiting symbiosis, where both organisms derive equal benefit from one another (Table 1).

**Fig 3.**
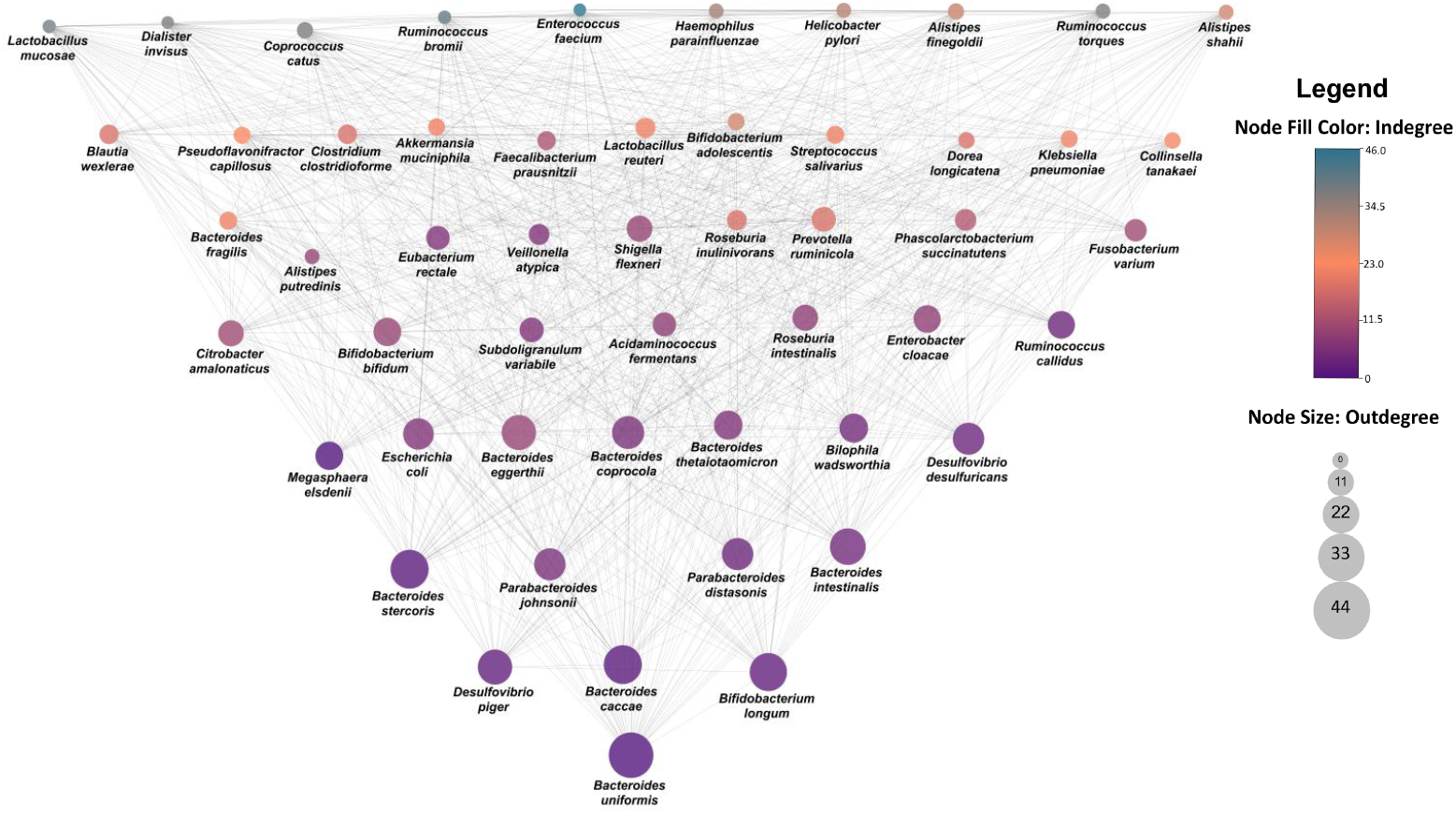
Association network constructed based on MSI values-. Association network constructed using the 793 common set of edges retained in HF and DMEM. The size of the node is proportional to it’s outdegree. The colour of the node is based on the indegree and varies according to the scale given on the right.

**Table 1.** Pairwise combinations of organisms that derive equal benefit from one another.

| Organism 1 | Organism 2 | MSI Value (%) |
| --- | --- | --- |
| *Alistipes finegoldii* | *Lactobacillus reuteri* | 0.4 |
| *Bacteroides thetaiotaomicron* | *Bacteroides intestinalis* | 3.5 |
| *Bacteroides uniformis* | *Bacteroides stercoris* | 1.6 |
| *Bifidobacterium bifidum* | *Bifidobacterium adolescentis* | 1.7 |
| *Collinsella tanakaei* | *Ruminococcus torques* | 0.9 |
| *Faecalibacterium prausnitzii* | *Ruminococcus torques* | 1.5 |
| *Haemophilus parainfluenzae* | *Enterococcus faecium* | 0.3 |
| *Pseudoflavonifractor capillosus* | *Lactobacillus reuteri* | 0.4 |
| *Ruminococcus torques* | *Prevotella ruminicola* | 1.1 |

In order to determine if there were few organisms that act as central nodes for interactions, we computed the network parameters. It was interesting to note that *Bifidobacterium bifidum* and *Ruminococcus torques* have the highest betweenness centrality **(Supplementary Table 5)**. These organisms have previously been demonstrated to degrade mucins, a characteristic not found in other *Bacteroides* species [48]. Hence, these organisms could act as key players to facilitate the interactions between other gut organisms. Moreover, we found that the average clustering coefficient of the association network drops from 0.488 on DMEM to 0.42 on HF. The characteristic path length also increased from 1.757 (DMEM) to 1.987 (HF). These observations indicate that the organisms tend to interact better on a nutritionally poor medium (DMEM) compared to a richer medium (HF). These observations corroborate those reported in §3.1.

### 3.4 Synergistic interactions between gut bacteria enhance amino acid biosynthesis

Amino acids have been demonstrated to play a vital role, both for the host and microbe. It has been previously reported that amino acids are predominantly secreted by the micro-organisms residing in the gut [49]. Moreover, these amino acids (along with other metabolites) act as a “communication medium” between the organisms [31,50]. Indeed, the amino acid biosyntheses in gut microbiomes are highly optimised to support cooperation between the organisms [31]. Hence, any improvement in the amino acid production capabilities of the organisms through metabolic exchanges can promote stronger co-operative interactions. Thus, to identify the enrichment in amino acid synthesising capabilities of the organisms, we first determined the number of amino acids synthesised by individual organisms and compared it with that from the joint metabolic networks. This analysis was carried out only on minimal glucose medium, since DMEM and HF medium contain all the amino acids.

From the dotplot **(Figure 4)**, we observed that the amino acid synthesising abilities were improved only for 18 organisms. In particular, it was striking to note that species belonging to *Lactobacillus* genus, especially *L. mucosae* and *L. reuteri* showed an enriched amino acid synthesising ability in the presence of many other gut organisms on a minimal glucose medium. Particularly, we observed that L-glutamate, L-aspartate, L-arginine and L-histidine were commonly synthesised by *L. mucosae* in association with other organisms **(Supplementary Table 6)**. This indicates an enhanced ability of these *Lactobacillus* spp. to work along with other organisms to produce vital metabolites – thereby pointing to their application in pro-biotics.

**Fig 4.**
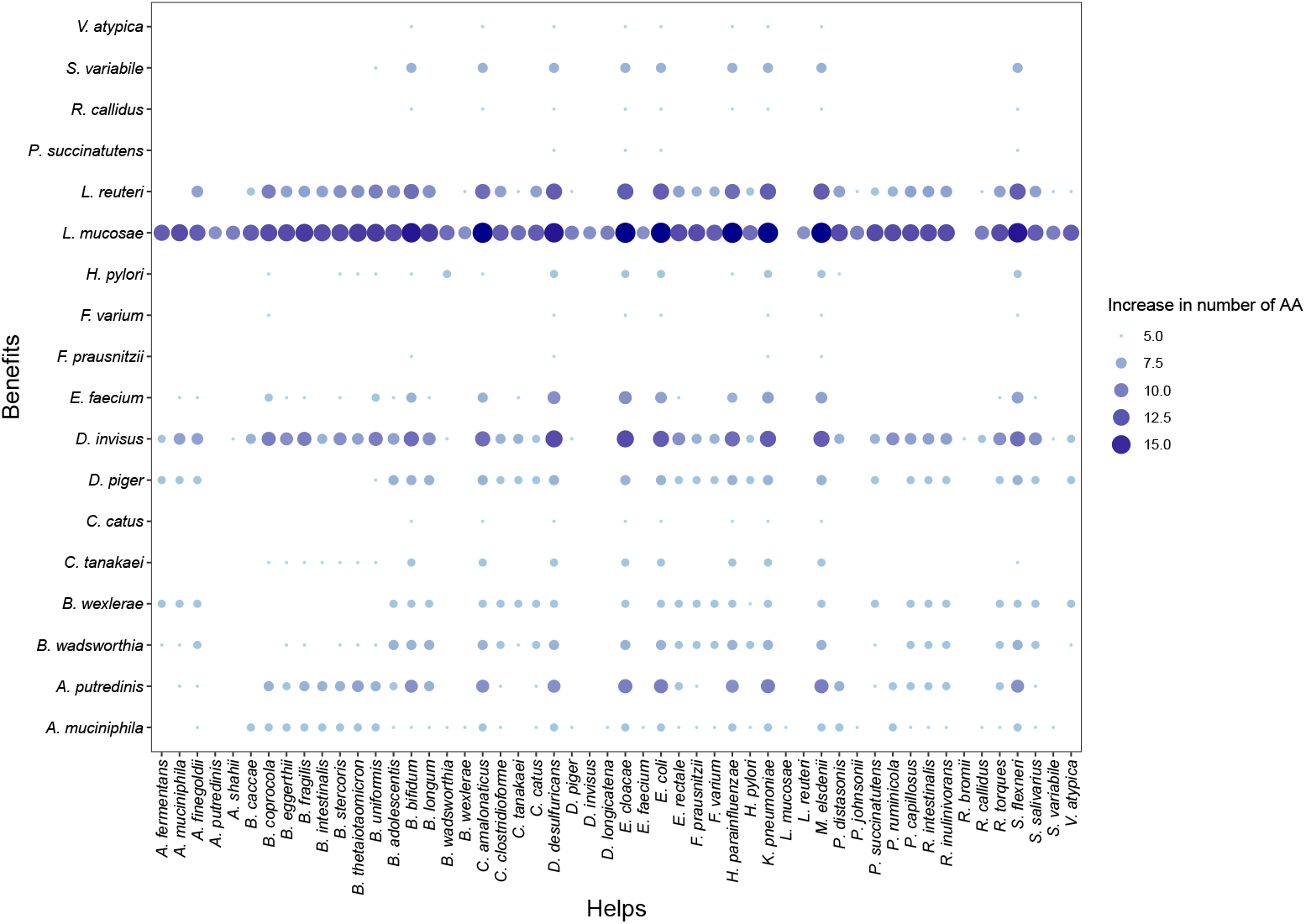
Dotplot showing the increase in amino acids synthesised by gut organisms. Dots show the increase in the number of amino acids synthesised by the organism on Y-axis in the presence of another organism (along X-axis). The size and colour of dots indicate the number of amino acids synthesised additionally in a pairwise interaction.

### 3.5 Pathway analyses reveal several metabolic exchanges in gut

Metabolic exchanges between the organisms in the gut were identified by systematically analysing all the pathways spanning the two organisms on both the medium conditions. We observed that the interdependencies between the organisms are highly contingent on the medium conditions. From our analyses, we found that the overall metabolic exchanges between the organisms reduced when the conditions were changed from DMEM to high fibre **(Figure 5)**. This observation reaffirms the intuition that the necessity for the organisms to depend on each other reduces in nutrient-rich conditions.

**Fig 5.**
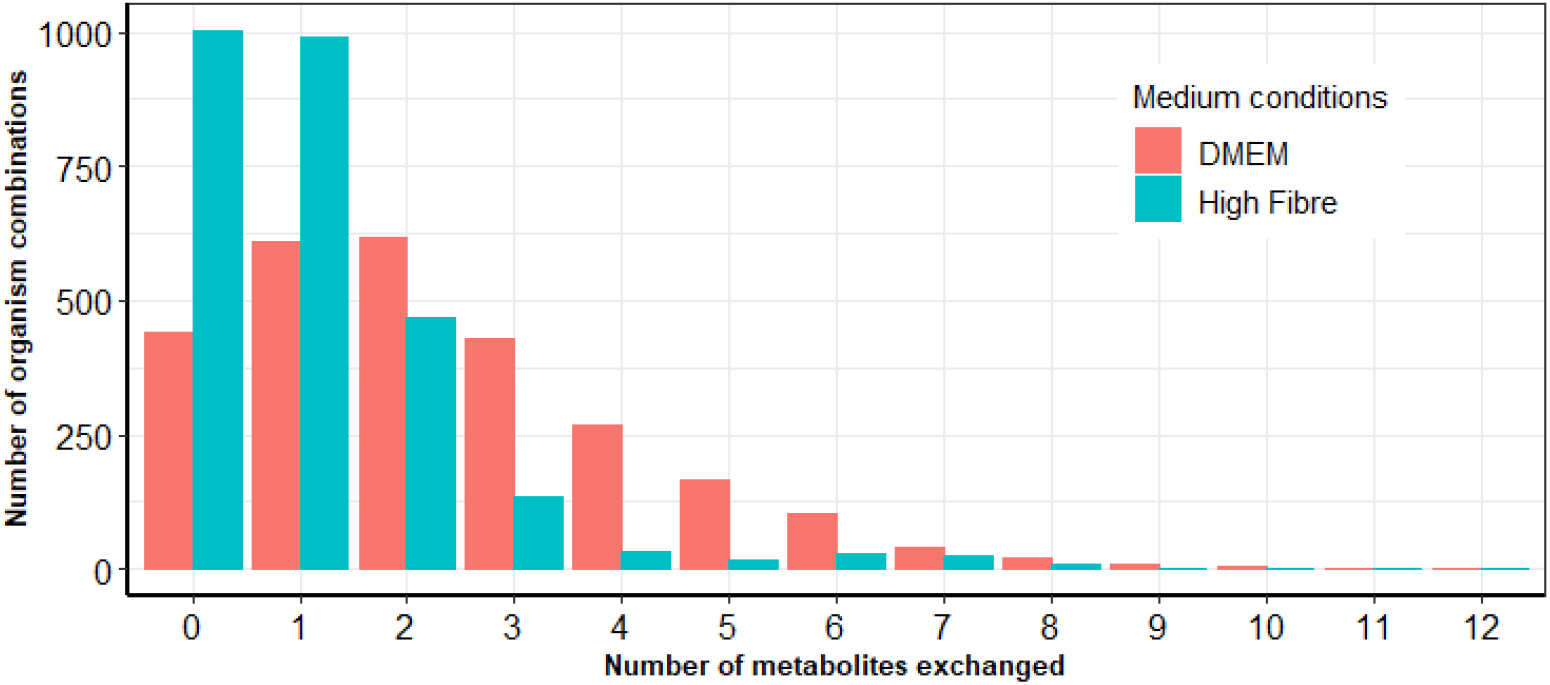
Barplot of metabolic exchanges. Barplot showing the number of metabolites exchanged on X-axis and the number of organism pairs exchanging metabolites on Y-axis. Two different conditions DMEM and HF are depicted. Note that the number of metabolites exchanged reduced in HF conditions compared to DMEM. Since metabolic exchanges are asymmetrical, note that there are 2704 entries (per condition) in this plot.

To test this observation, we scrutinised every pair of organisms and probed into the metabolic exchanges. Overall, we found that the maximum number of metabolites exchanged reduced from 12 (DMEM) to 8 (HF). Interestingly, we noted that the metabolic exchange between few pairs of organisms had increased in HF compared to that found in DMEM. Particularly, we found that *Ruminococcus bromii, Bifidobacterium bifidum* and *Bilophila wadsworthia* receive several metabolites from the other gut organisms in HF **(Figure 6)** in contrast to that on DMEM **(Supplementary Figure S4)**. We noted that many of these exchange metabolites either pertained to the mucin-degrading metabolic products or SCFAs **(Supplementary Table 7)**. For instance, we observed that many of the metabolites from mucin degradation pathway in *Ruminococcus bromii* exchanged with *Bifidobacterium bifidum* were involved in either cell wall synthesis or energy metabolism.

**Fig 6.**
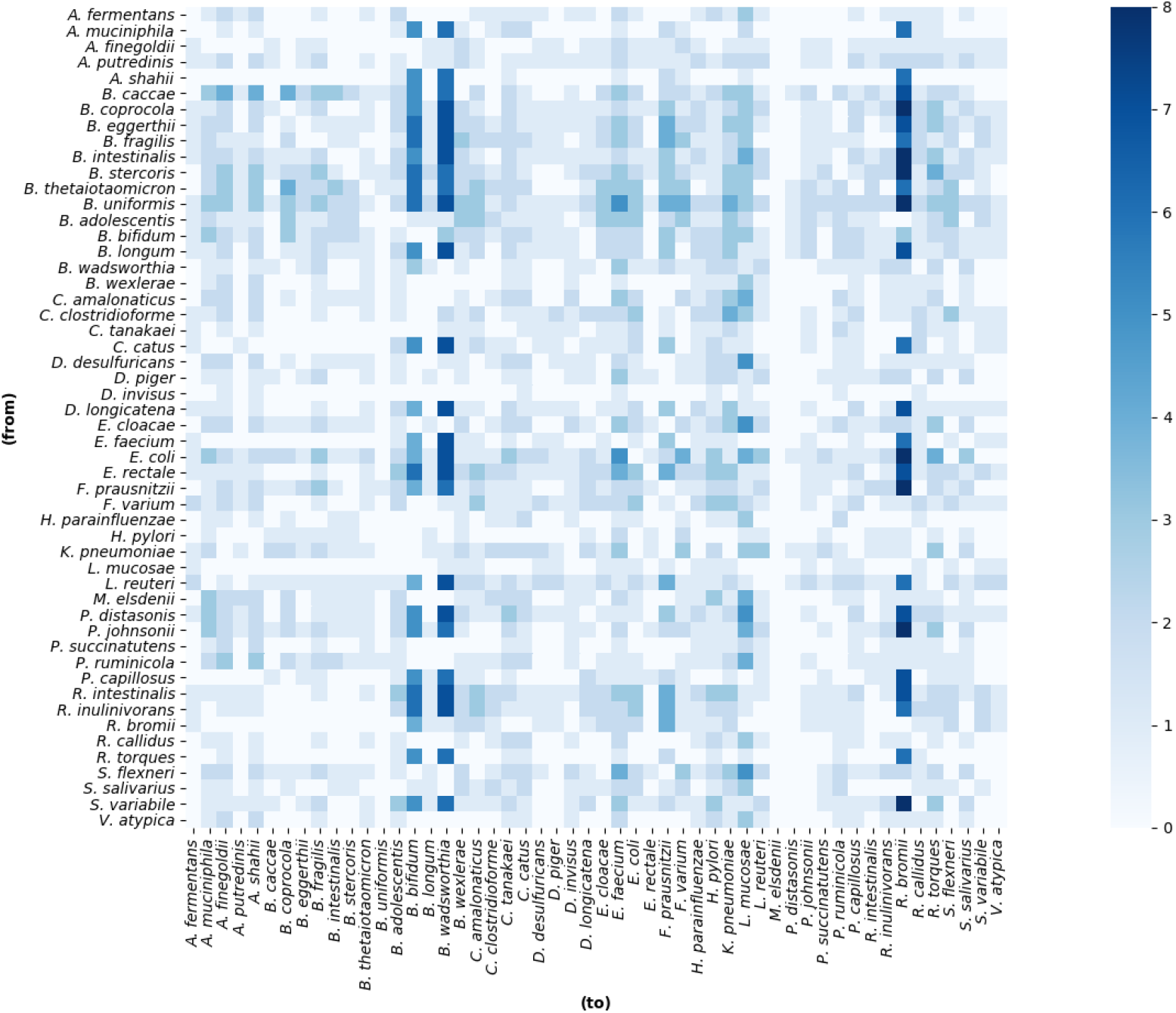
Heatmap showing metabolic exchanges on HF. The number of unique metabolites exchanged between the organisms is shown. X-axis represents the organisms to which metabolites are transferred, Y-axis represents the organisms from which metabolites are transferred. Note that a few organisms such as *Ruminococcus bromii* and *Bilophila wadsworthia* receive many metabolites from several organisms in the gut.

**Fig 7.**
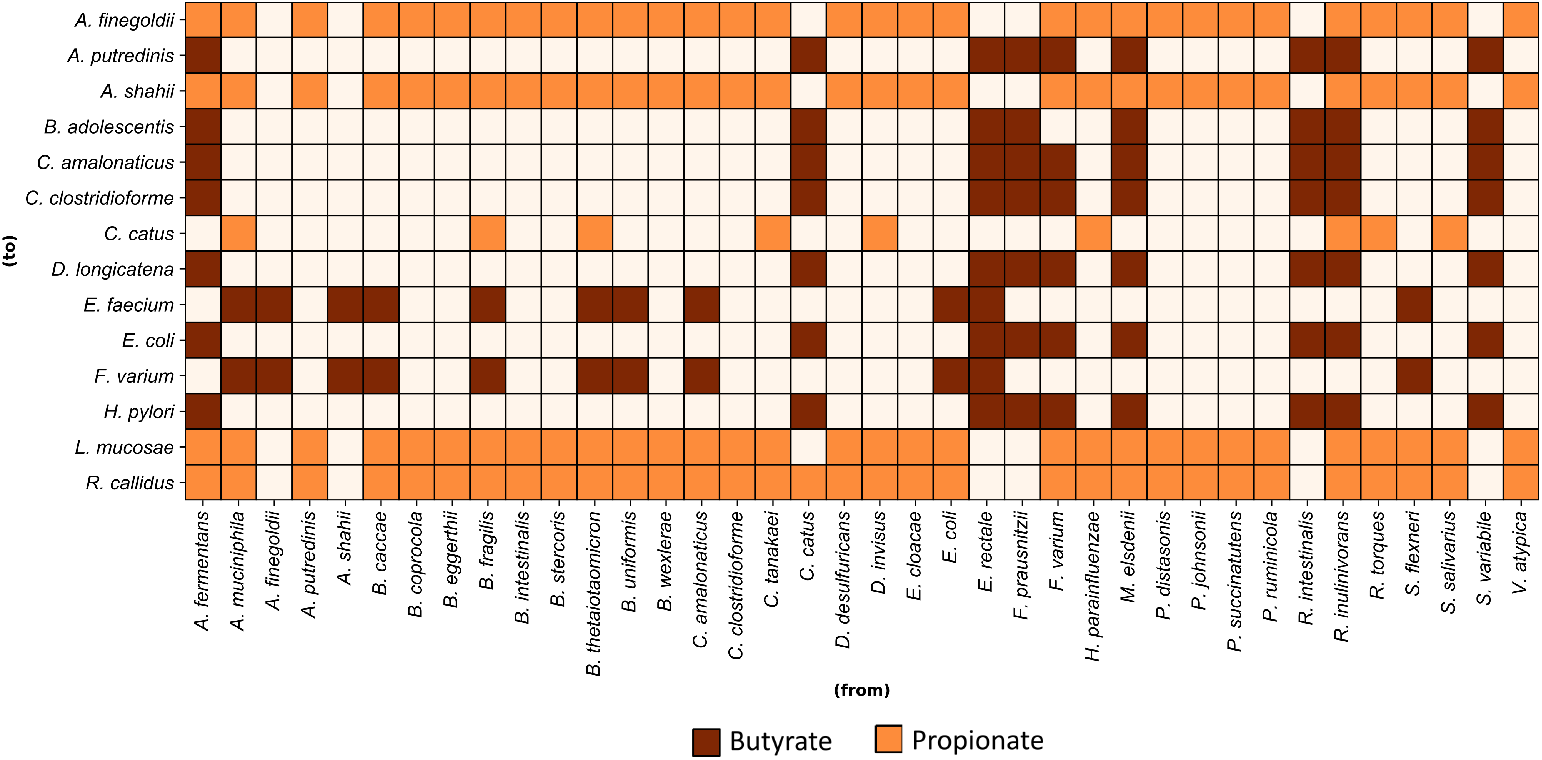
SCFAs exchange profile. SCFAs: butyrate indicated in maroon and propionate indicated in orange are transferred from organisms on Y-axis to the organisms on X-axis. Note that several organisms belonging to Bacteroides exchange propionate, while the classic butyrate producers are involved in butyrate exchange.

Further, the metabolic exchanges on DMEM medium conditions, revealed that the organisms predominantly exchange vitamins and amino acids such as nicotinic acid, L-ornithine, nucleotides including thymidine (nucleotide metabolism) and spermidine, and various small molecules involved in central carbon metabolism, such as acetate, glycerol, lactate, 2-keto butryric acid and citrate. On HF conditions, we found that many of the exchange metabolites were derived from mucin metabolism, and these predominantly played a role in cell wall biosynthesis. All these metabolites are available in **Supplementary Table 8**.

We also identified the common pool of metabolites which are exchanged between the organisms in both these conditions. Interestingly, we observed that this pool comprised mainly of propionate and butyrate, which are SCFAs well known to have impact on several functions of human body [51,52]. It was striking to note the *F. prausnitzii, A. fermentans, R. inulinivorans*, which are known butyrate producers in the gut [53–55], were primarily involved in exchanging this metabolite with many other gut inhabitants. Further, we also found that several species belonging to Bacteroidetes, such as *B. uniformis, B. eggerthii* and *A. putredinis* exchange propionate.

## 4 Discussion

Microbes in the gut possess astounding capabilities and carry out a wide array of functions. Within the gut microbiome, the organisms exhibit different types of interactions. Investigating these interactions plays crucial roles in deciphering the mechanisms and understanding the roles of the organisms in the microbiome. In this study, we chose 52 most commonly occurring gut microbes and applied a novel graph-theoretic framework to unravel the microbial interdependencies.

Our results from these analyses are four-fold. Firstly, based on the MSI values on two different medium conditions based on increasing richness - DMEM and HF, we observe few phyla such as Firmicutes, Fusobacteria, and Proteobacter exhibited significantly higher MSI values in DMEM compared to HF. In addition, we observed that microbial cooperation is higher in DMEM medium compared to HF. This is as expected, since the organisms, when grown in a gut-like rich media have access to many more nutrients compared to the DMEM conditions. Due to this, the interdependencies between the organisms is bound to reduce. These observations are corroborated well by previous experiments, where it was shown that minimal medium conditions often promote the organisms to cooperate and complement their biosynthetic capabilities [10,56,57].

Secondly, we studied the influence of phylogenetic and metabolic distance on interactions between the micro-organisms. On DMEM, the phylogenetic distance showed a weak positive correlation with MSI; however, no such correlations were observed in HF conditions. Such a difference in the correlations could be due to the metabolic support offered by the specific metabolites produced by the respective phylum. Further, we observed that the metabolic distance does not show significant correlations with MSI. This observation is in agreement with the results previously reported [58]. Such a non-significant correlation between MSI and metabolic distance also indicates that the metabolites shared between the two gut organisms can be easily synthesised (primary metabolites) as opposed to the requirement from secondary metabolism. This observation also aligns well with those reported previously [59]. We also found that *Eubacterium rectale* and *Roseburia intestinalis* offer a better support in the HF conditions compared to DMEM, which could be due to the enrichment of these species in high-fibre food sources, as reported elsewhere [60].

Thirdly, based on the association networks derived from metabolic dependencies, we observed that the microbes in the gut exhibit a trophic-level arrangement—a few organisms such as *B. uniformis, B. stercoris, B. thetaiotamicron and B. caccae*, irrespective of the medium conditions, serve as ‘helpers’ and support many other organisms in the gut. These ‘helper’ organisms have been reported to have broader metabolic capabilities, including a repertoire of enzymes to breakdown complex carbohydrates [61]. These factors, along with the association network **(Figure 3)**, suggest a spatial distribution of these organisms in the gut depending on their metabolic capabilities, as also shown in previous studies [62]. Further, due to their central role in the network, these species can play a critical role in community assembly [63].

Next, we show that synergistic interactions between the gut bacteria promote the ability of amino acid biosynthesis. Particularly, we noted that microbial interactions of lactic acid bacteria with other gut microbes helped in the production of several amino acids such as L-glutamate, L-aspartate and L-arginine. Although lactic acid bacteria have already been studied and explored as probiotics [64], combinations with other gut organisms may prove to be more effective.

Finally, from the pathway analyses, we note for many pairs of organisms, that the number of metabolites exchanged decreases with increasing richness of medium conditions. A few pairs of organisms involving *R. bromii and B. bifidum* receive more metabolites in the HF condition compared to DMEM, which were mainly from mucin degradation pathway. Further, specific metabolite analyses showed that many of these metabolites exchanged between the organisms were a part of energy metabolism, nucleotide and vitamin biosynthesis. In addition, we also observed the exchange of SCFAs butyrate and propionate, whose roles have been elucidated in several previous studies [51,52,65]. These results indicate that SCFAs can prove to be effective pre-biotics, promoting interactions between critical gut organisms.

Our study does have some limitations. First, our algorithm provides a static snapshot of the interactions happening in the gut microbiome. Nevertheless, graph-based approaches to study microbial communities are highly useful and present a complementary picture to constraint-based analyses. Further, while some of our results have been confirmed with previously reported experiments, we also present many results that remain to be validated. While these provide a fertile ground for future research, many challenges remain, particularly in the context of understanding and characterising the metabolome, or tracking metabolite exchanges in complex microbial communities. Finally, our predictions are also heavily contingent on the quality and the level of curation of the metabolic models, notably the transport and exchange reactions in the models, which can heavily influence our predictions.

Overall, in this study, we have presented a novel modelling framework to carry out a systematic analysis of the gut microbiome. Using this framework, we demonstrated the microbial interactions in the gut and predicted several putative metabolic exchanges. These predictions provide mechanistic insights and testable hypotheses to guide experimental validations. It is possible to come up with a number of strategies to perturb and study these networks, on the basis of the simulations discussed herein. For instance, it may be possible to modulate microbial communities through the perturbation of key metabolic exchanges. Further, it may also be interesting to understand the changes in microbial interactions during alterations in the gut caused due to a variety of factors such as antibiotic usage and inflammatory bowel disease (IBD). Notably, the techniques discussed here are generic and thus can be extended for analysing metabolic interactions in any type of microbial community.

## Supporting information captions

**Supplementary Figure S1 – Correlations between MSI and phylogenetic distance on DMEM.** Each dot represents an organism pair. The trendline is indicated in blue color. Spearman rank value for this correlation *ρ* = 0.22, *p* =1.4 × 10^−15^

**Supplementary Figure S2 – Correlations between MSI and phylogenetic distance on HF.** Each dot represents an organism pair. The trendline is indicated in blue color. No significant Spearman rank value for this correlation was observed.

**Supplementary Figure S3 – Correlations between Metabolic distance and phylogenetic distance.** Each dot represents an organism pair. The trendline is indicated in blue colour. Spearman rank value for this correlation *ρ* = 0.19, *p* = 7.66 × 10^−11^

**Supplementary Figure S4 - Heatmap showing metabolic exchanges on DMEM diet conditions.** The number of unique metabolites exchanged between the organisms is shown. X-axis represents the organisms to which metabolites are transferred, Y-axis represents the organisms from which metabolites are transferred.

**Supplementary Table 1** - List of 52 organisms used in this study (CSV)

**Supplementary Table 2** - Constituents of the three environmental conditions used in this study (Minimal glucose, DMEM, HF) (CSV)

**Supplementary Table 3** - Organism pairs grouped into three categories and their MSI values on DMEM and HF conditions (CSV)

**Supplementary Table 4** - List of 52 organisms along with the Number of organisms ‘helped’ and the organisms from which benefits are derived on the two different medium conditions (DMEM, HF) (XLSX)

**Supplementary Table 5** - Betweenness centrality values of 52 organisms (XLSX)

**Supplementary Table 6** - A 52 × 52 matrix with the amino acids synthesised by every organism (rows) in the presence of the other (columns) (XLSX)

**Supplementary Table 7** - A 52 × 52 matrix with the unique metabolites exchanged from one organism (rows) to the other (columns) in HF conditions (XLSX)

**Supplementary Table 8** - A list of metabolites and the number of times it is exchanged with the other (XLSX)

## Acknowledgements

A.R. acknowledges fellowship from the Initiative for Biological Systems Engineering, IIT Madras.

