## Supplementary Figures for "Unraveling microbial interactions in the gut microbiome"

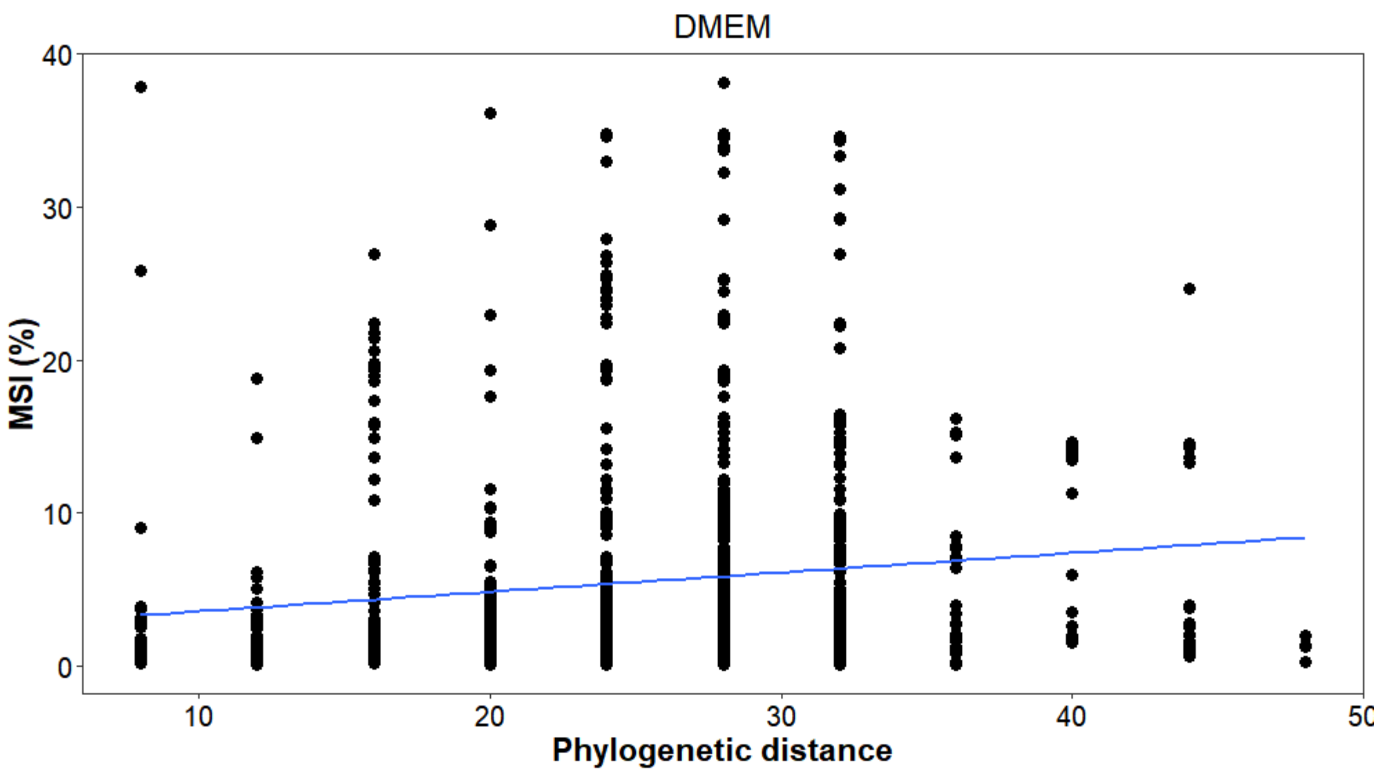


**Supplementary Figure S1 – Correlations between MSI and phylogenetic distance on DMEM**. Each dot represents an organism pair. The trendline is indicated in blue color. Spearman rank value for this correlation $\rho = 0.22$, P =$1.4*10^{-15}$

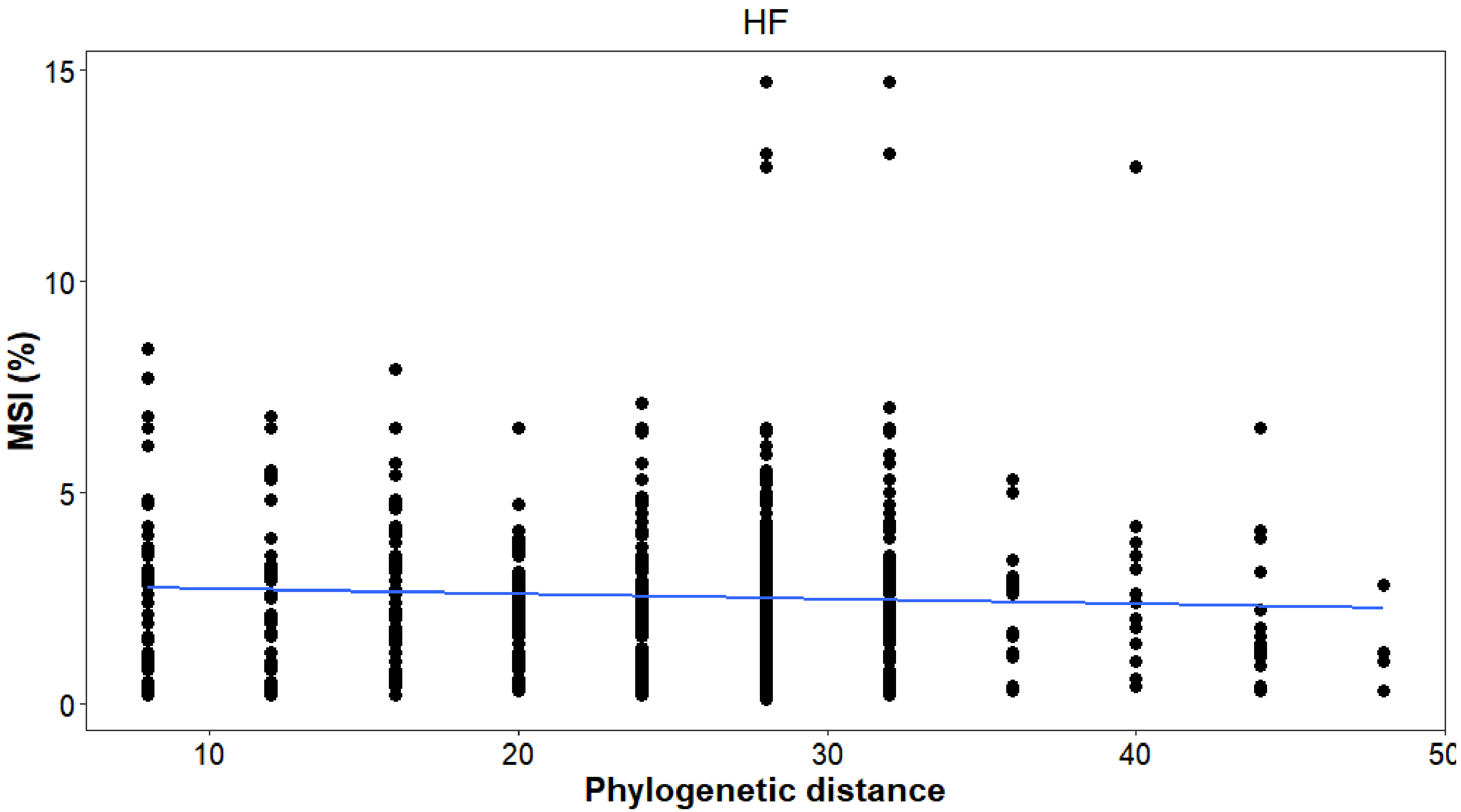


**Supplementary Figure S2 – Correlations between MSI and phylogenetic distance on HF**. Each dot represents an organism pair. The trendline is indicated in blue color. No significant spearman rank value for this correlation was observed.

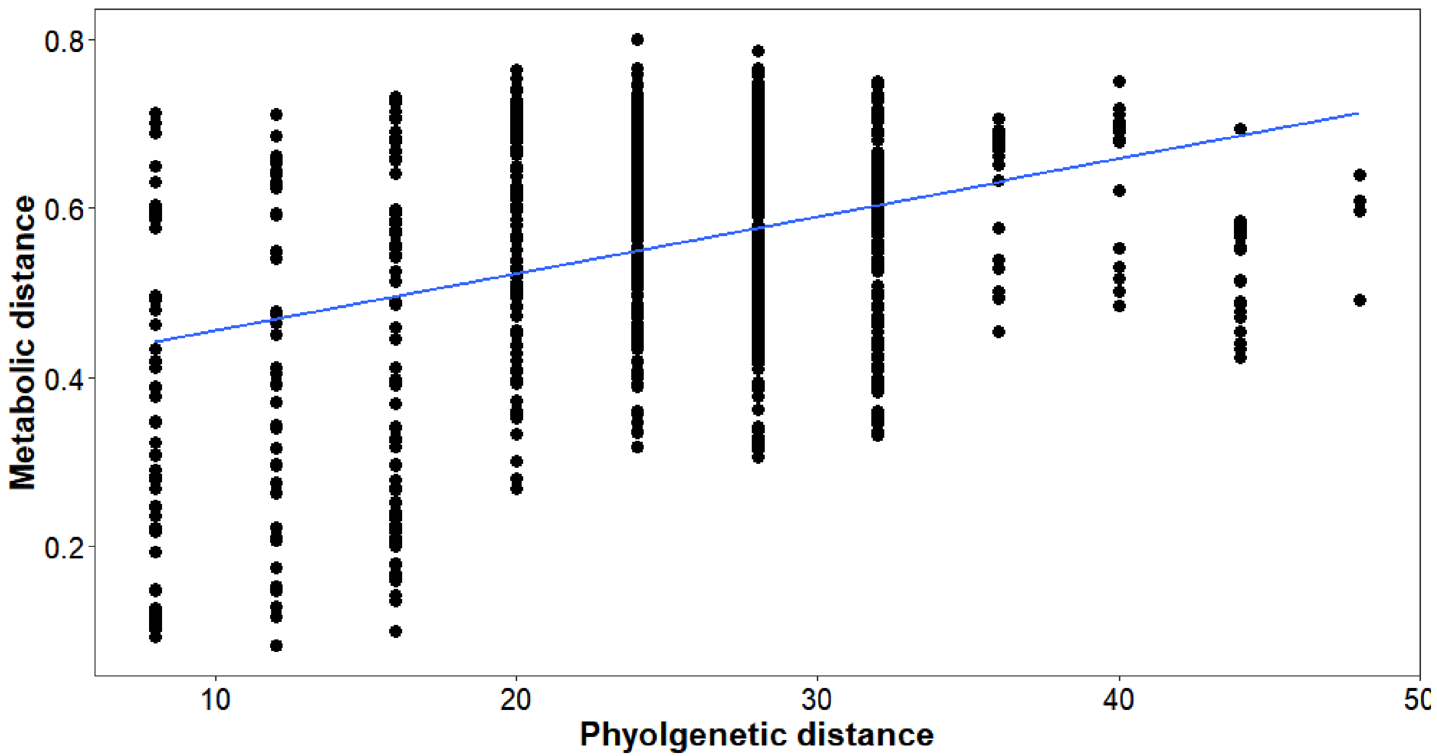


**Supplementary Figure S3 – Correlations between Metabolic distance and phylogenetic distance**. Each dot represents an organism pair. The trendline is indicated in blue color. Spearman rank value for this correlation $\rho = 0.19$, P =$7.66*10^{-11}$

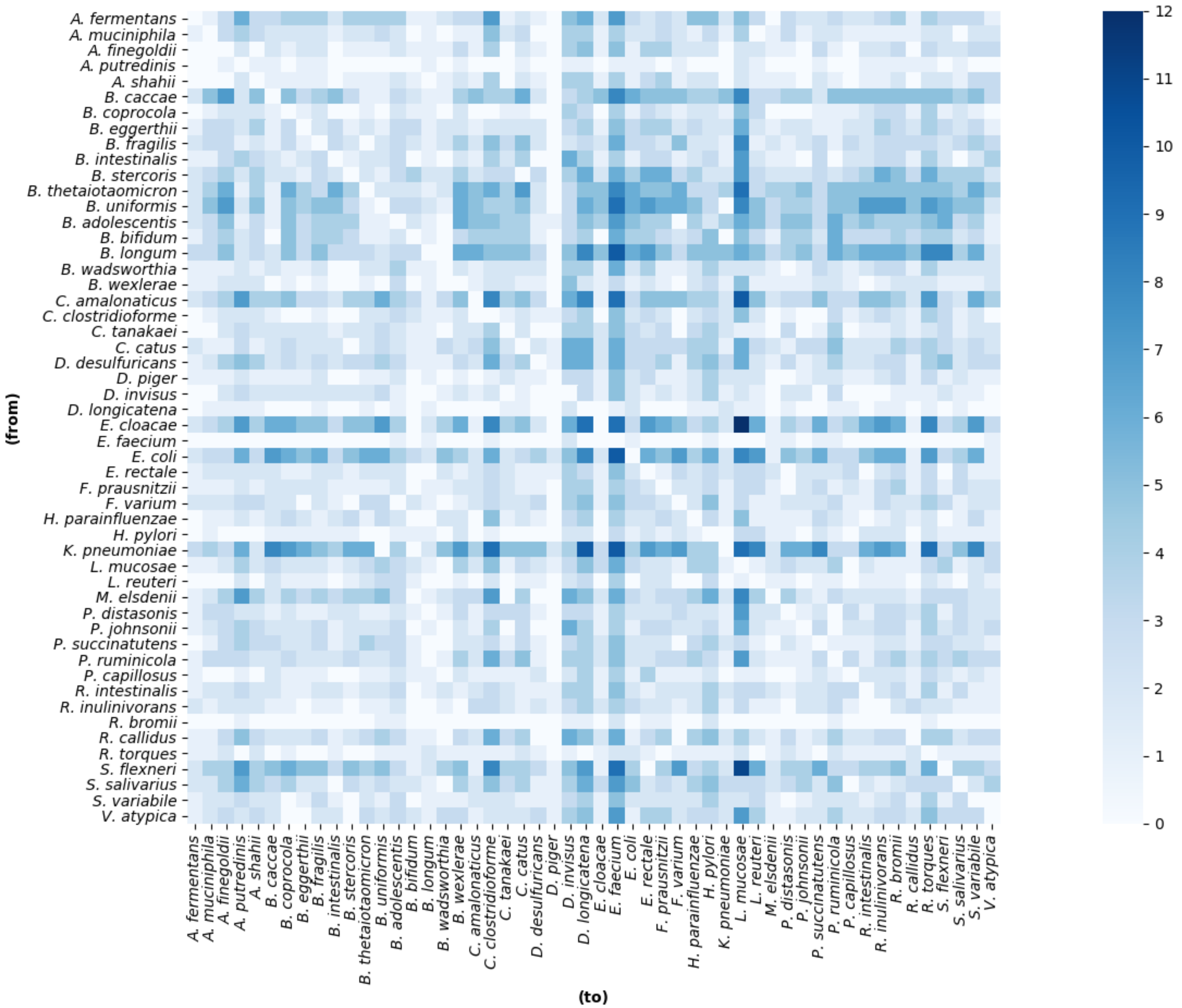


**Supplementary Figure S4 Heatmap showing metabolic exchanges on DMEM diet conditions.** The number of unique metabolites exchanged between the organisms is shown. X-axis represents the organisms to which metabolites are transferred, Y-axis represents the organisms from which metabolites are transferred.
